# “Dual protease type XIII/pepsin digestion offers superior resolution and overlap for the analysis of histone tails by HX-MS”

**DOI:** 10.1101/788463

**Authors:** James Mullahoo, Terry Zhang, Karl Clauser, Steven A. Carr, Jacob D. Jaffe, Malvina Papanastasiou

## Abstract

The N-terminal regions of histone proteins (tails) are dynamic elements that protrude from the nucleosome and are involved in many aspects of chromatin organization. Their epigenetic role is well-established, and post-translational modifications (PTMs) present on these regions contribute to transcriptional regulation. While hydrogen/deuterium exchange mass spectrometry (HX-MS) is well-suited for the analysis of dynamic structures, it has seldom been employed to analyze histones due to the poor N-terminal coverage obtained using pepsin. Here, we test the applicability of a dual protease type XIII/pepsin digestion column to this class of proteins. We optimize online digestion conditions using the H4 monomer, and extend the method to the analysis of histones in monomeric states and nucleosome core particles (NCPs). We show that the dual protease column generates many short and overlapping N-terminal peptides. We evaluate our method by performing hydrogen exchange experiments of NCPs for different time points and present full coverage of the tails at excellent resolution. We further employ electron transfer dissociation (ETD) and showcase an unprecedented degree of overlap across multiple peptides that is several fold higher than previously reported methods. The method we report here may be readily applied to the HX-MS investigation of histone dynamics and to the footprints of histone binding proteins on nucleosomes.

---

Histones are highly conserved proteins found primarily in the nucleus that organize into higher order structures with DNA to form nucleosomes, the basic building block of chromatin. Nucleosomes are composed of ∼147 bp of DNA wrapped around a histone octamer, which is assembled by two H2A/H2B dimers, and an H3/H4 tetramer [1]. The highly basic and largely flexible histone N-termini (tails) extend out of the nucleosome and are readily accessible to enzymes that can add (writers) or remove (erasers) post-translational modifications (PTMs). These PTMs may act in a synergistic or sequential fashion to recruit various histone binding proteins (readers) and remodelers, affecting downstream transcriptional processes [2] and chromatin structure [3]. Nucleosome core particles that include histones and DNA can be reconstituted *in vitro*, and serve as a useful model system for the study of chromatin biology.

Despite their critical function, the three dimensional structure of the tails and their role in mediating intra- and inter-nucleosome interactions with both DNA and proteins of the transcriptional machinery are largely unexplored. Since the first crystal structure of the octameric histone core was determined [4], a number of high resolution histone (and variant) structures have been reported in complexes with chaperones, nucleosomes and nucleosomal arrays (622 entities in 378 PDB entries, rcsb.org) [5]; in most of these structures, parts of the histone tails could not be resolved due to their high flexibility. Peptides corresponding to segments of the tails and modified versions of those have been employed as an alternative approach and have been successfully resolved in complexes (∼270 entities). Analytical methods that will tackle these challenges and will allow for the structural analysis of full histone sequences in a physiological context are highly desirable.

Hydrogen/deuterium exchange-mass spectrometry (HX-MS) is a powerful method for probing highly dynamic regions of proteins (such as the N-terminal regions of histones) that may be inaccessible to other methods [6]. In the few published studies that have attempted to probe histone dynamics, the tails were represented poorly, either by the lack of peptides in this region or by the identification of long peptides (∼50 amino acids; for a detailed background see [7]). In all these studies, a traditional bottom-up approach was employed, where proteins were digested rapidly into peptides using pepsin, a non-specific endoprotease that is functional under HX-MS quenching conditions. Neprosin, a selective prolyl endoprotease, was reported to improve resolution of the H3 and H4 tails compared to pepsin [8], but has not yet been used for monitoring in-solution histone dynamics. More recently, we improved coverage and redundancy of H2A, H3 and H4 tails further (peptides ∼20-25 amino acids long) through the use of Cathepsin-L and showcased its applicability to the analysis of the in-solution dynamics of H3 and H4 in a monomeric context [7]. Sequence resolution of H3 and H4 was significantly increased using Cathepsin-L followed by on-line pepsin digestion, but background signals also increased, necessitating additional washes with high concentrations of guanidine hydrochloride between runs to maintain a low background. This is a known issue with offline, in-solution digestions where a high enzyme:substrate ratio is required, therefore immobilized protease columns are highly preferred.

Recently, a dual protease type XIII/pepsin column became commercially available and it is slowly being adopted for in HX studies [9, 10]. Protease type XIII from *Aspergillus saitoi* was introduced in the HX-MS field in 2003 [11] and has mainly been used for the digestion of proteins in-solution [12–15], although an immobilized version used for deuterium exchange studies of proteins has also been described [16, 17]. Like pepsin, it is a non-specific enzyme and maintains its activity in the presence of reducing and denaturing agents [15] commonly required in HX-MS experiments. Its cleavage preferences at the C-terminal side of basic amino acids [14] are complementary to pepsin, enabling improved peptide maps with a high percentage of overlapping fragments [9, 10, 16]. Here, we introduce the protease type XIII for studying histone tails by HX-MS. We optimize online digestion conditions using the dual protease type XIII/pepsin column and showcase excellent resolution for all core histones either at the MS1 or MS2 level.

## Experimental Section

### Materials

Human recombinant histones (H2A; M2502S, H2B.1; M2505S, H3.1; M2503S and H4; M2504S) were purchased from New England BioLabs Inc (Ipswich, MA). Mononucleosomes, recombinant human (16-0009) containing H2A (P04908), H2B (O60814), H3.1 (P68431), and H4 (P62805) were purchased from EpiCypher (Durham, NC). D; 151882). All other reagents were from Sigma-Aldrich (St. Louis, MO).

### Sample preparation

Individual histones (5 uL of 0.2 ug/uL in guanidine HCl, pH 2.5) or mononucleosomes (10 uL of 0.2 ug/uL in guanidine HCl, pH 2.5) were injected to the MS via an automated HDx-3 PAL™ system (LEAP Technologies, Morrisville, NC). Samples were digested online for 2 min at 200 uL/min using an immobilized protease type XIII/pepsin column (w/w, 1:1, NBA2014002, NovaBioAssays, Woburn, MA), operated at 8 °C and were trapped onto an Acclaim™ PepMap™ 300 µ-Precolumn™ (C18, 1 × 15 mm, 163593, Thermo Fisher Scientific, Waltham, MA) using solvent A (0.1% formic acid (v/v)). Peptides were separated onto a Hypersil Gold C18 (1 × 50 mm, 1.9 uM, 25002-051030, Thermo Scientific, Waltham, MA) at 40 uL/min using solvents A and B (0.1% v/v formic acid in acetonitrile). The following gradient was applied: 3% B to 10% B in 0.1 min, to 35% B in 6 min, to 95% B in 0.1 min; kept at 97% B for 1.9 min and returned to initial conditions in 0.9 min. Both the trap and analytical column were operated at 0 °C. For hydrogen exchange experiments, mononucleosomes (4 uL of 0.4 ug/uL) were mixed with ice-chilled deuterated PBS (36 uL, final D_2_O content during reaction 90% v/v). Samples were quenched at 60, 600 and 3600 s, with 6 M guanidine HCl (20 uL) to a final pH 2.2 (0.6% formic acid). Samples were injected as described above and incubated in the loop for 1 min prior to online digestion. Full deuteration controls were prepared by overnight labeling at 37 °C using an Eppendorf™ Thermomixer™ R (Thermo Scientific, Rockford, IL) and injected as described above.

### MS data acquisition

MS analysis was carried out on an Orbitrap Fusion™ Lumos™ Tribrid™ Mass Spectrometer (Thermo Fisher Scientific) using a spray voltage of 3.5 kV, capillary temperature 220 °C and vaporizer temperature 50 °C. Full MS scans were acquired in the *m/z* range 270-1500, with an AGC target 5e5 and 60,000 resolution (at *m/z* 200). Peptides with charge states 2 to 8 were selected for MS/MS fragmentation using ETD or HCD. Parameters were as follows: AGC target 4e5, loop count 12 and isolation window 3 *m/z*. ETD reaction times were 50 ms for charges 2-3, 25 ms for charge 4 and 20 ms for charges 5-8; HCD collision energy was 28% for all charges. Data were acquired in profile mode and peptides were identified using Spectrum Mill Proteomics Workbench (prerelease version B.06.01.202, Agilent Technologies). Searches were performed using ESI QExactive in the Instrument menu. The fragmentation mode was ETD only for ETD experiments and All for HCD experiments. A non-specific enzyme search was performed using a fasta file containing sequences of human histones, pepsin and protease type XIII. Peptide and fragment tolerances were at ±20 ppm and peptide FDR at 1%. Further processing, manual validation of the data and deuterium uptake measurements occurred in HDExaminer (Sierra Analytics).

## RESULTS AND DISCUSSION

### Optimization of digestion conditions

The proteolytic activity of the dual protease column was optimized using the H4 monomer followed by ETD fragmentation [18]. Due to the hydrophilic nature of the tails, we tested varying digestion times (1-3 min) and flow rates (100-300 μL/min) aiming to obtain high N-terminus coverage and repeated measurements of amino acids from overlapping peptides (often referred to as redundancy). In all conditions, spectra corresponding to intact H4 were not detected, indicating that digestion was complete. The number of tail peptides identified (up to amino acid 30) ranged from 6-11 (Fig. 1A) with the total digestion volume being the largest contributor; it was found that 400-600 μL provided optimum number of peptides and intensities. Using less than 400 μL was insufficient to remove guanidine HCl, resulting in decreased peptide intensities, while using digestion volumes above 600 μL resulted in the loss of shorter peptides during trapping. In this optimal range, increasing the pressure (from 80 bar to 255 bar) by increasing the flow rate (from 100 μL/min to 300 μL/min) had minimal effect in the number of peptides identified, we therefore chose the minimum optimum digestion volume (2 min at 200 μL/min) to prevent back exchange.

**Figure 1.**
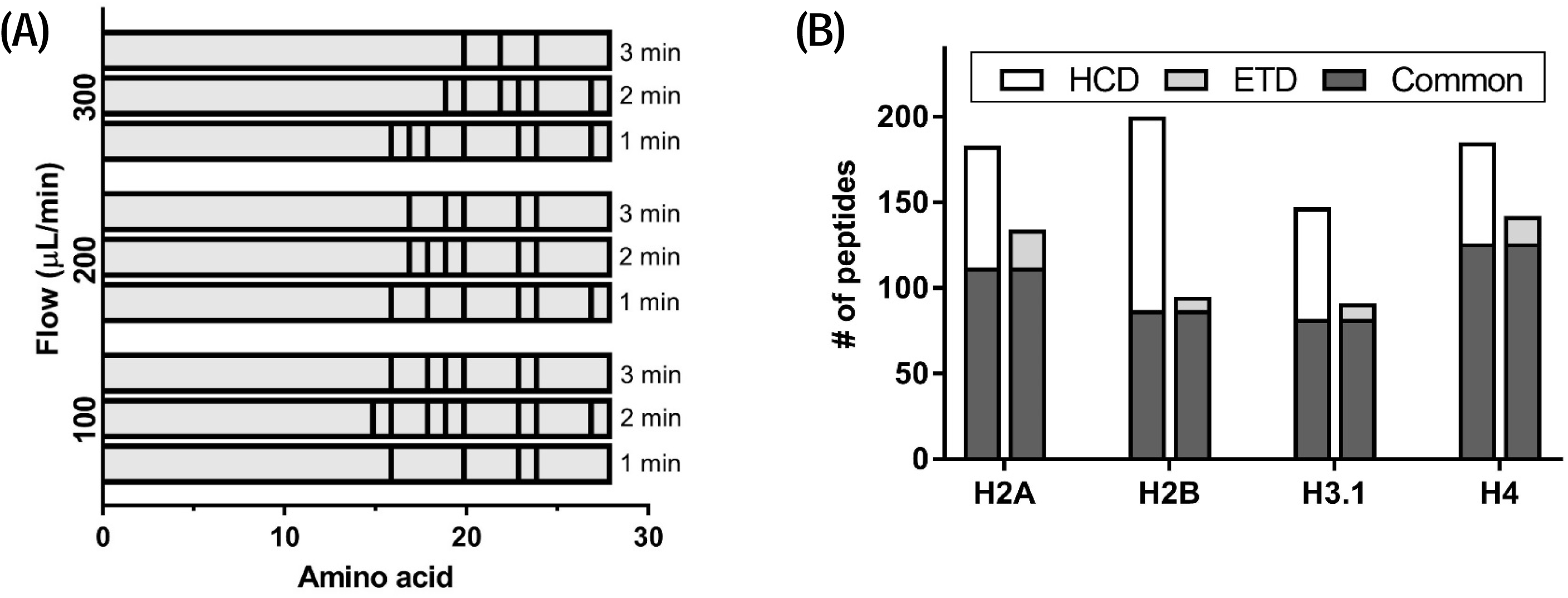
Optimization of digestion conditions and peptide identifications. (A) Effect of flow rate and digestion time on peptide identification for the H4 N-terminus (amino acids 1-30) using ETD. Only peptides that start at amino acid 1 are shown. Black lines indicate the end amino acid of each peptide identified. (B) Total number of unique peptides identified by HCD (white) and ETD (light grey) at optimized digestion conditions (2 min at 200 μL/min). Common peptides are shown in dark grey.

### Histone peptide identifications by HCD and ETD

We then tested the overall protein coverage and reproducibility of the column using the four core histones and performed digestions in triplicates. To generate a comprehensive map of the peptides produced upon digestion with the dual protease column, we applied both HCD that is available in Orbitrap systems and ETD fragmentation that is better suited for the analysis of high charge state cationic molecules [19] and is anticipated to provide better resolution of tail peptides. The number of peptides identified ranged from ∼90 to ∼200 with a 42% (H2B) to 63% (H4) overlap between the two methods (Fig. 1B). HCD resulted in a greater sequence depth compared to ETD, due to the slower scan rate of the latter (∼⅔ the rate of HCD) that results in less frequent peptide sampling [20]. Both methods however resulted in complete coverage of H2B and H4 and >94% for H2A and H3.1 (Fig. 2A, 2B). Despite the larger pool of peptides identified by HCD and the higher repeated AA overlap, most tail peptides were identified by both HCD and ETD, rendering either method suitable for HX measurements of histone tails at the MS1 level. H3.1 peptides. Pearson correlation coefficients of peptide intensities obtained in at least two replicates were >0.90. Standard deviations using HCD were inferior compared to ETD, presumably due to the higher number of small peptides identified (Fig. S1A, Table S1). These belonged mainly to the histone folds (starting at AA ∼30 for H2A, H2B, H4 and AA 46 for H3 and extending to the C-terminus) and could still be qualified for HX measurements, however caution should be exercised during their selection due to their reduced reproducibility. Profiling the specificity of the column using these peptide identifications revealed cleavage sites characteristic to pepsin (Leu, Met) and protease type XIII (Arg) activities [14] (Fig. S1B). As histones form the stable core of nucleosomes, we next attempted to determine peptides generated for nucleosomal core particles (NCPs) by a standard in-line digestion using the dual protease column. Peptides generated upon digestion with the dual protease column were profiled at the MS1 level, as would typically occur for HX-MS measurements. Using the peptide libraries generated above, we extracted the precursor ions and inspected them in HDExaminer. For all histones, the number of peptides validated ranged from ∼117 (H2A) to ∼178 (H4) (Fig. 2C). Compared to peptides identified from monomeric histones, identifications from the nucleosome matched with >85% of peptides from H3.1 and H4 monomers, ∼78% from H2B, and ∼60% from H2A. While the total number of peptide identification decreased, sequence coverage was unaffected, remaining similar between NCPs and monomers. Interestingly, the number of peptides corresponding to the tails was only marginally reduced relative to the monomers, yielding similar coverage. Taken together, these results indicate that despite the increase in sample complexity (histone octamer ∼110 kDa), the presence of DNA and the lower amount of protein injected (0.5 μg per histone in NCPs vs. 1 μg in monomers), the developed method generated high resolution peptide maps for both the tails and the histone cores.

**Figure 2.**
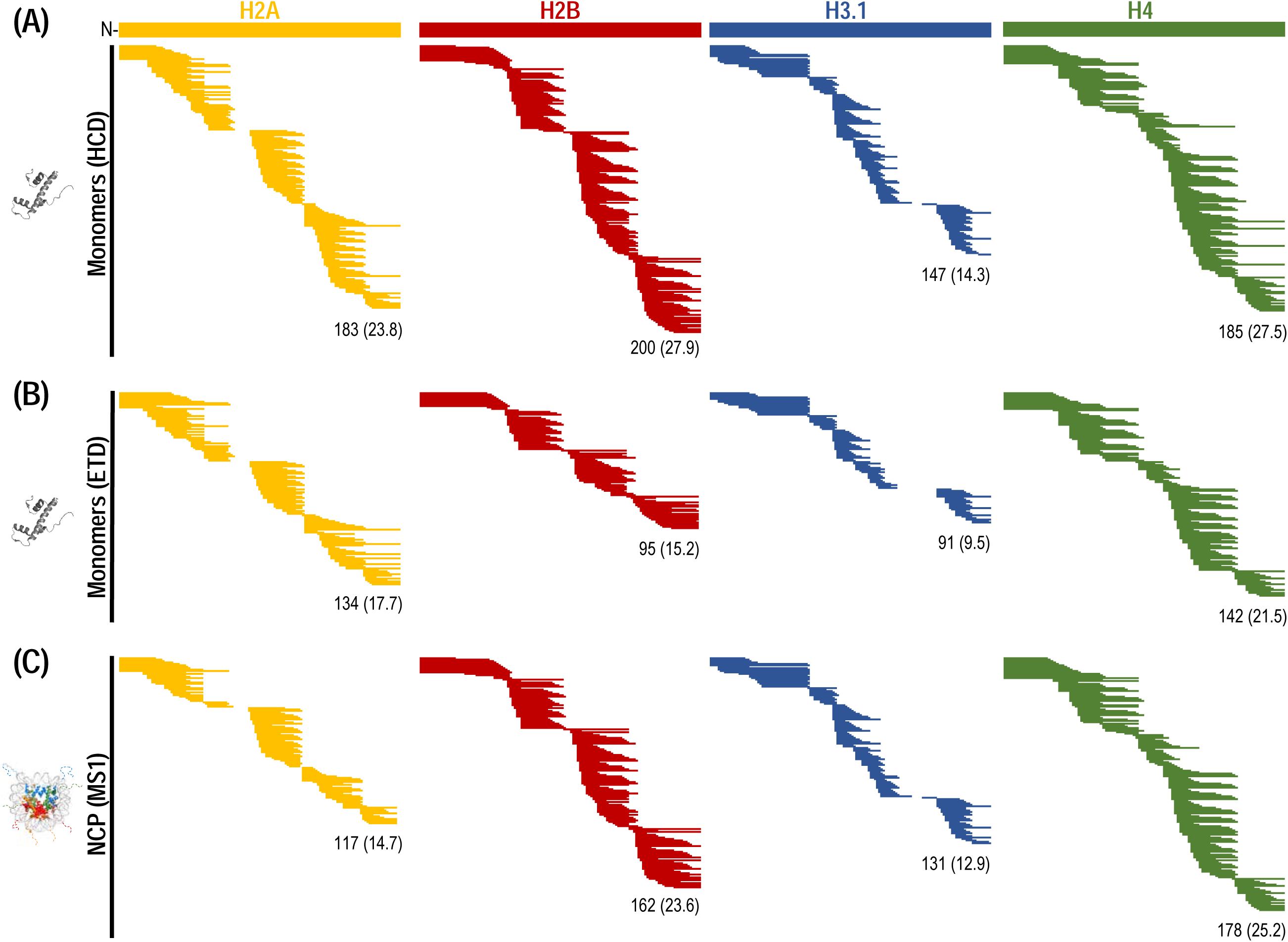
Peptide maps generated upon digestion with the dual protease column. Coverage maps of histone monomers using (A) HCD and (B) ETD. Each line corresponds to a unique peptide identified in at least two replicates (n=3). (C) Peptides identified in mononucleosomes at the MS1 level. The total number of peptides identified is given at the bottom of each figure along with the average AA overlap value (parenthesis). Peptide identifications are given in Table S1.

Given the importance of the tails and their PTMs in regulating transcriptional proCesses, we next foCused our analysis on the improvements the dual protease column offers towards for deuterium measurements at the MS1 level that is the most commonly used approach in deuterium exchange studies. We also calculated repeated AA overlap at the residue level, corresponding to the number of times a given AA has been identified in detected peptides (Fig. S2A); higher AA overlap values are desirable since they provide higher structural resolution and further validation of deuterium measurements. For all histones, a large number of overlapping tail (first 30 AAs) peptides with overhanging residues at their C-termini (e.g. H4) or both termini (e.g. H3) were detected upon HCD fragmentation (Fig. 2A, Table S1), well outperforming peptide identifications reported to date for this class of proteins using other proteases. For example, pepsin generates only one large peptide (1-50) for H3.1, whereas cathepsin-L generates two shorter peptides (1-21, 1-22) in addition to 1-50, improving resolution in this region [7]). In contrast, the dual protease column generated up to 16 overlapping peptides for H3.1 in residues 1-30 and resulted in almost complete residue resolution for AAs 5-8 and 15-26, covering 70% of the tail (Fig. S2B). Similarly, near-single residue resolution was obtained for AAs 14-28 (50% of the tail) in H2A and 15-26 (43% of the tail) in H4. These are areas that contain known target residues for PTMs and the ability to calculate exchange rates at the residue level using a traditional bottom-up approach opens new avenues for histone structural studies. An increase in the site-resolution may further be feasible through the use of advanced computational approaches, using overlapping peptides that do not share common termini [21]. Single residue resolution was achieved in mononucleosomes, with the exception of H2A that was slightly lower (AAs 17-28). AA overlap values calculated *per* residue were in the range of 6-35 with the majority exceeding a value of 10 (Fig. S2B). These correspond to ∼2 to 4-fold increase compared to previous studies with pepsin or Cathepsin-L/pepsin [7, 22]. Similar redundancy values have been reported for other, non-histone proteins using the dual protease digestion scheme, even for proteins up to 900 residues long [17], indicating the unique advantages the dual protease digestion has to offer. Next, we were interested to assess the method we developed on deuterated samples to evaluate how the presence of deuterium that results in isotopic expansion will impact protein coverage and redundancy. We employed the dual protease digestion configuration and performed deuterium exchange measurements of NCPs for three different time points (60 s, 600 s, 3600 s). In total, we were able to analyze 66 peptides in H2A (80% coverage), 73 in H2B (97% coverage), 51 in H3 (80% coverage) and 70 in H4 (96% coverage) for their D_2_O content. Between 40-60% of the undeuterated peptides previously identified under regular H2O buffer conditions (Fig. 2C) were able to be re-identified and analyzed in their deuterated states. This decrease is attributed to the limited incubation time with guanidine HCl (1 min) prior to digestion, in contrast to the undeuterated ones that were fully denatured. Despite the overall decrease in peptides generated, the tails were represented by a high number of peptides, offering unprecedented redundancy in this region compared to previously reported methods. We detected 19 peptides in H2A, 9 in H2B, 15 in H3 and 24 in H4; this corresponded to a marginal decrease in average redundancy values (8 to 12) compared to the denatured, undeuterated peptides (11 to 17) (Fig. S2C). This is anticipated, as flanking histone tails extend outside the boundaries of NCPs and are accessible for proteolysis. The deuterium uptake for all histones is depicted in the form of a heat map for individual time points (Fig. 3). It is shown that the tails are almost fully deuterated over the time course of the experiment in contrast to the histone folds that exhibit low deuterium uptake, even at the latest time point. For the investigation of protein dynamics, a higher labeling temperature or longer labeling time points is highly recommended.

**Figure 3:**
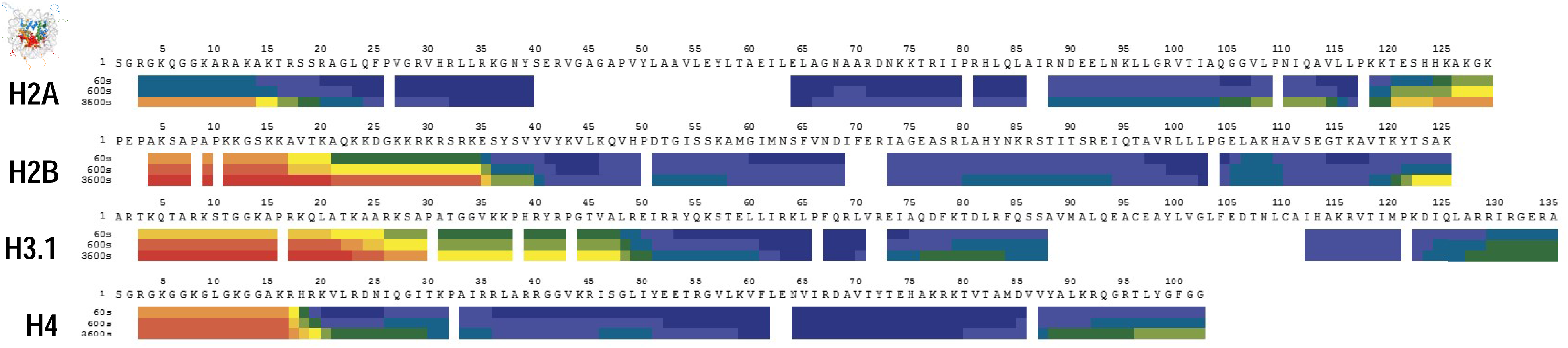
Deuterium uptake of histones in NCPs by HX-MS. Heat maps depicting H-to-D exchange for each histone in nucleosome core particles (NCPs) (A). Amino acid sequences for individual histones are shown on top. Color coding of heat maps is based on deuterium update (%) calculated based on normalizing D-uptake values to a fully deuterated NCP control sample as described in Mayne, 2016. Individual time points of exchange are 60, 600 and 3600 seconds. Individual D-uptake values are given in Table S1.

### Extending resolution of histone tails to the residue level

Localization of the deuterium content at the residue level occurs either at the peptide level through the identification of many overlapping peptides with overhanging N- and C-terminal residues [21] or using ECD/ETD (MS2) that prevents hydrogen scrambling [18]. We further took advantage of ETD capabilities to obtain residue resolution maps across the extensively modified tail regions (first 30 AAs). The majority of tail peptides identified (92%) were detected with charges +4 to +8 that correlated with length, rendering them excellent candidates for ETD fragmentation (Fig. S3). We calculated overlap *per* residue at the MS2 level, considering fragment ions from overlapping peptides and different charge states that result in separate MS2 spectra (Fig. S2A). Pairing complementary fragment ions from overlapping peptides in this is anticipated to yield much richer ion ladders and separate true N- and C-terminal ions from noise [20]. Summing all cleavage sites from overlapping peptides, and omitting residues N-terminal to proline at which ETD cleavage does not occur, full residue coverage of the tails was obtained, with the exception of H2B. Residue overlap was highest for H2A and H4 where >90% of the residues had values >10, followed by H3.1 and H2B where >60% had values >10 (Fig. 4). Despite the decreased coverage observed (∼80%) for H2B compared to the other histones, high overlap values were obtained for most lysine residues that undergo N-terminal modifications that are important.

**Figure 4.**
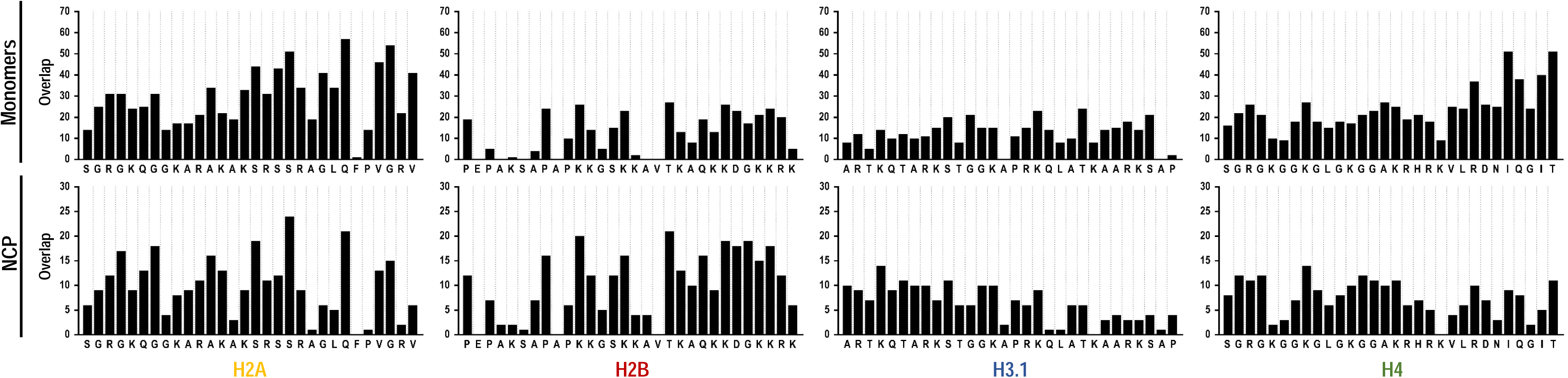
Resolution and amino acid overlap of histone tails at the MS2 level. AA overlap of histone tails (AAs 1-30) in monomers and NCPs fragmented by ETD at the MS2 level. These have been calculated as depicted in Fig. S2A.

We next analyzed whole NCPs using the same ETD workflow as for histone monomers. Pearson’s correlations of peptide intensities were >0.93 indicating excellent reproducibility (Fig. S1). The number of overlapping peptides identified within the tails was in the range of ∼8-11 for individual histones, providing full coverage in this region. Similar to the monomers, overlap values at the MS2 level were highest for H2A and H4 (80% of residues had values >5), followed by H2B (73%) and H3.1 (64%) (Fig. 4). To our knowledge, this is the first time that overlap values of this magnitude are reported for NCP histones at residue resolution. In a first of its kind, recent study, where ETD was employed for the analysis of NCPs following pepsin digestion, the resolution obtained for the first 30 AAs by single peptides (∼40-50 amino acids long), was ∼13% for H2A and H2B and ∼48% for H3.1 and H4 indicating many non-resolved segments [22]. Using the dual protease column and performing ETD on all peptides identified, we demonstrate here that >97% residue resolution is achieved in NCPs for H2A, H3.1 and H4, and >90% for H2B. Of note, these results are obtained using <25 msec ETD reaction time for charge states >+4. Further improvements in resolution and overlap may be achieved using longer reaction times or a higher amount of fluoranthene ions, however further optimization was beyond the scope of this study.

In conclusion, the dual protease digestion column shows great promise for the structural analysis of histone tails by HX-MS. The extensive number of overlapping peptides generated provides unique resolution of the tails at the MS1 level at unprecedented redundancy, circumventing the need for ETD fragmentation in studies where single residue resolution is not required. We demonstrate that our method is applicable to the analysis of histones in a mononucleosomal context. We further demonstrate that the use of ETD provides single residue resolution of the tails across their entire length at unprecedented redundancy. The methods presented herein are not restricted by sample complexity or buffer composition, creating new capabilities for histone studies addressing protein interactions and the role of PTMs in transcriptional processes and on chromatin dynamics.

## Acknowledgements

This work was supported by Thermo Scientific and by a grant from the National Institute of Health (NIH) to J.D.J. at the Broad Institute of MIT & Harvard (U54-HG008097). The authors would like to acknowledge Susan Klaeger (Broad Institute) for useful discussions on ETD methods, Susan Bird (Thermo Scientific) and Jeff Morrow (Sierra Analytics) for support.

## Data availability

The original mass spectra and sequence database have been deposited in the public proteomics repository MassIVE and are accessible at ftp://MSV000083879@massive.ucsd.edu when providing the dataset password: histone. If requested, also provide the username: MSV000083879. This dataset will be made public upon acceptance of the manuscript.

---

**Figure S1A.**
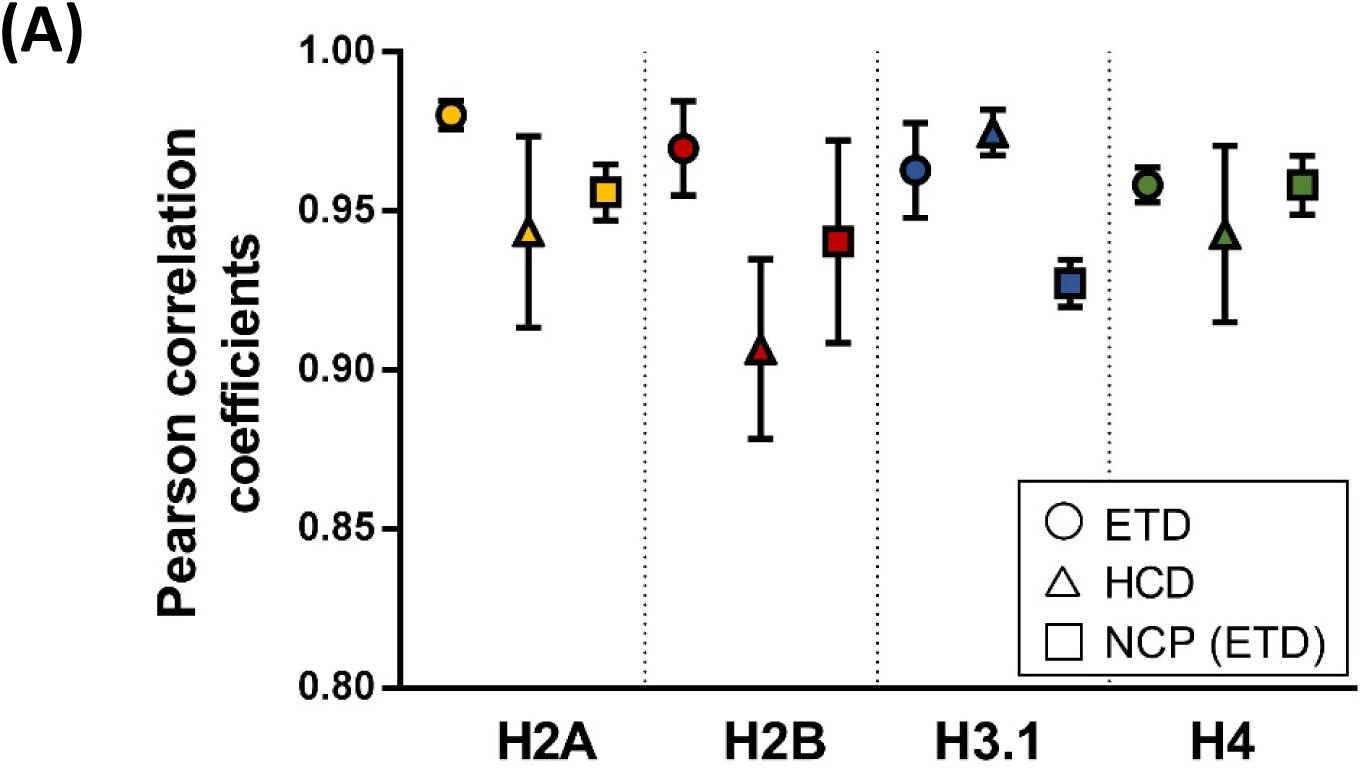
Reproducibility of the dual protease column derived from histone digestion using ETD and HCD. Pearson correlation coefficients were calculated using normalized peptide intensities obtained in at least two digestions (n=3) of histone monomers and mononucleosomes (NCP). Error bars represent standard deviations from the mean value.

**Figure S1B.**
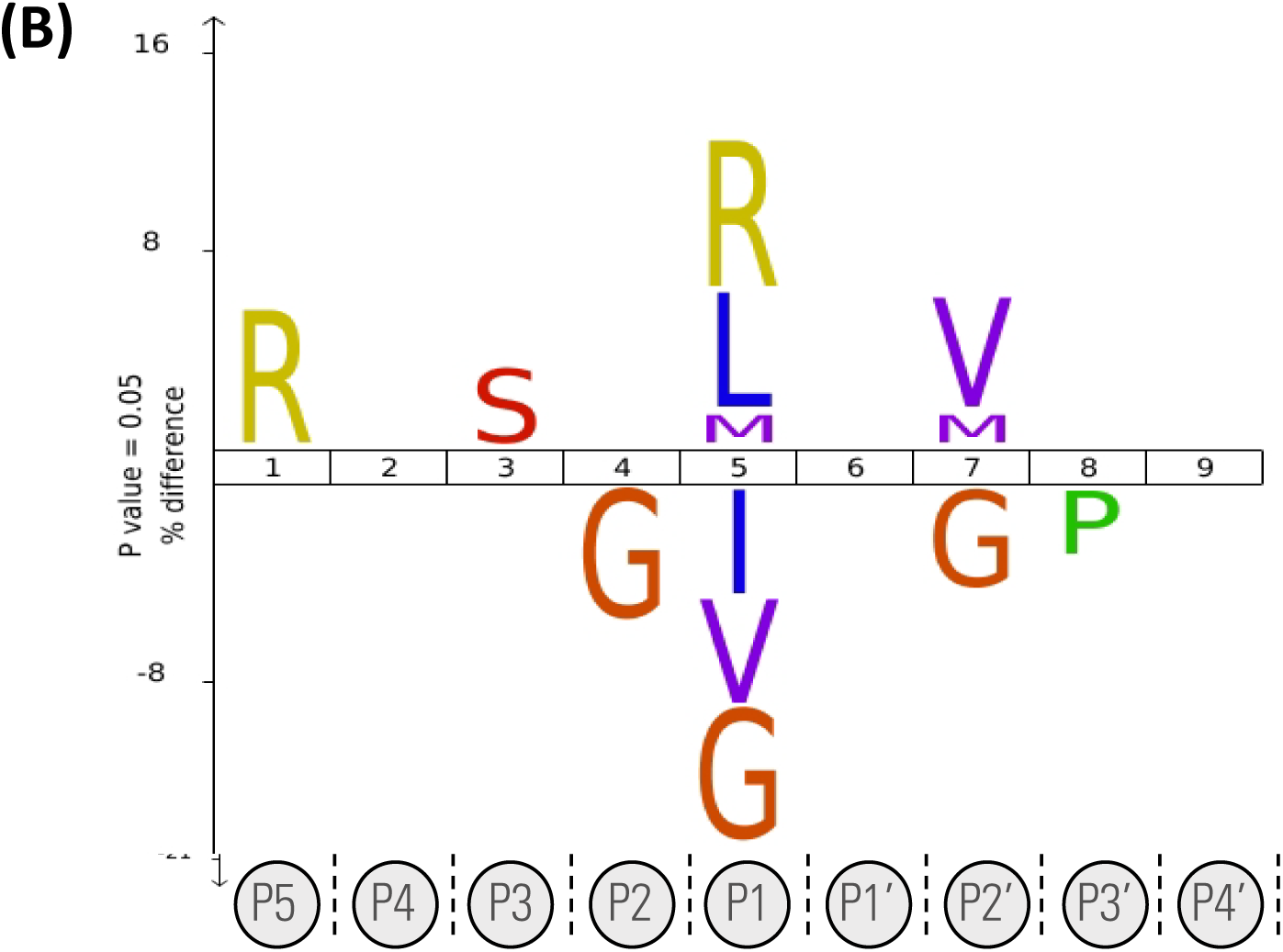
Specificity of the dual protease XIII/pepsin column derived from histone digestion at pH 2.5. To build the IceLogo (Colaert *et al*, 2009), we employed peptides identified from all four histones at quench conditions (200 uL/min for 2 min) and discarded redundant cleavage sites. Peptide sequences were aligned and loaded to the IceLogo web application. Peptides originating from histone sequences were used as a reference set. Over-represented amino acids around the cleavage site (P1) in the experimental set are indicated with positive values whereas underrepresented amino acids with negative values. Residues are colored based on their physicoChemical properties.

**Figure S2.**
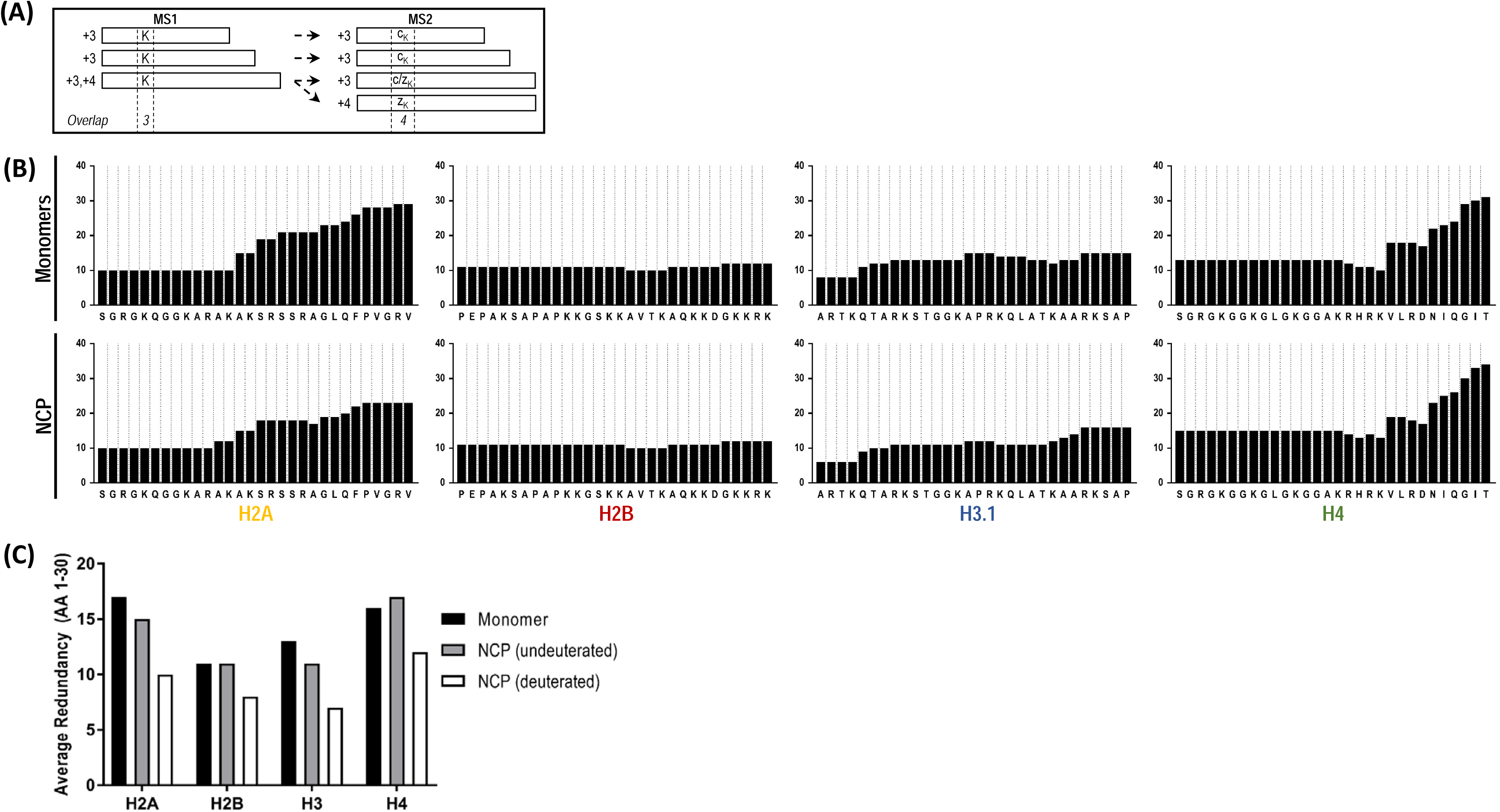
Overlap of histone tails at the MS1 level. **(A)** Cartoon representation of amino acid overlap calculations. At the MS1 level, overlap corresponds to the times a given AA has been identified in overlapping peptides (depicted in a rectangular shape). At the MS2 level, overlap corresponds to the number of times an AA has been identified from fragmentation spectra collected for any given charge state of a peptide. **(B)** Overlap values for AAs 1-30 of histone monomers and NCPs upon HCD identification at the MS1 level. **(C)** Average redundancy of the histone tails for AAs 1-30 in the monomers, NCPs (undeuterated) and NCPs (deuterated).

**Figure S3.**
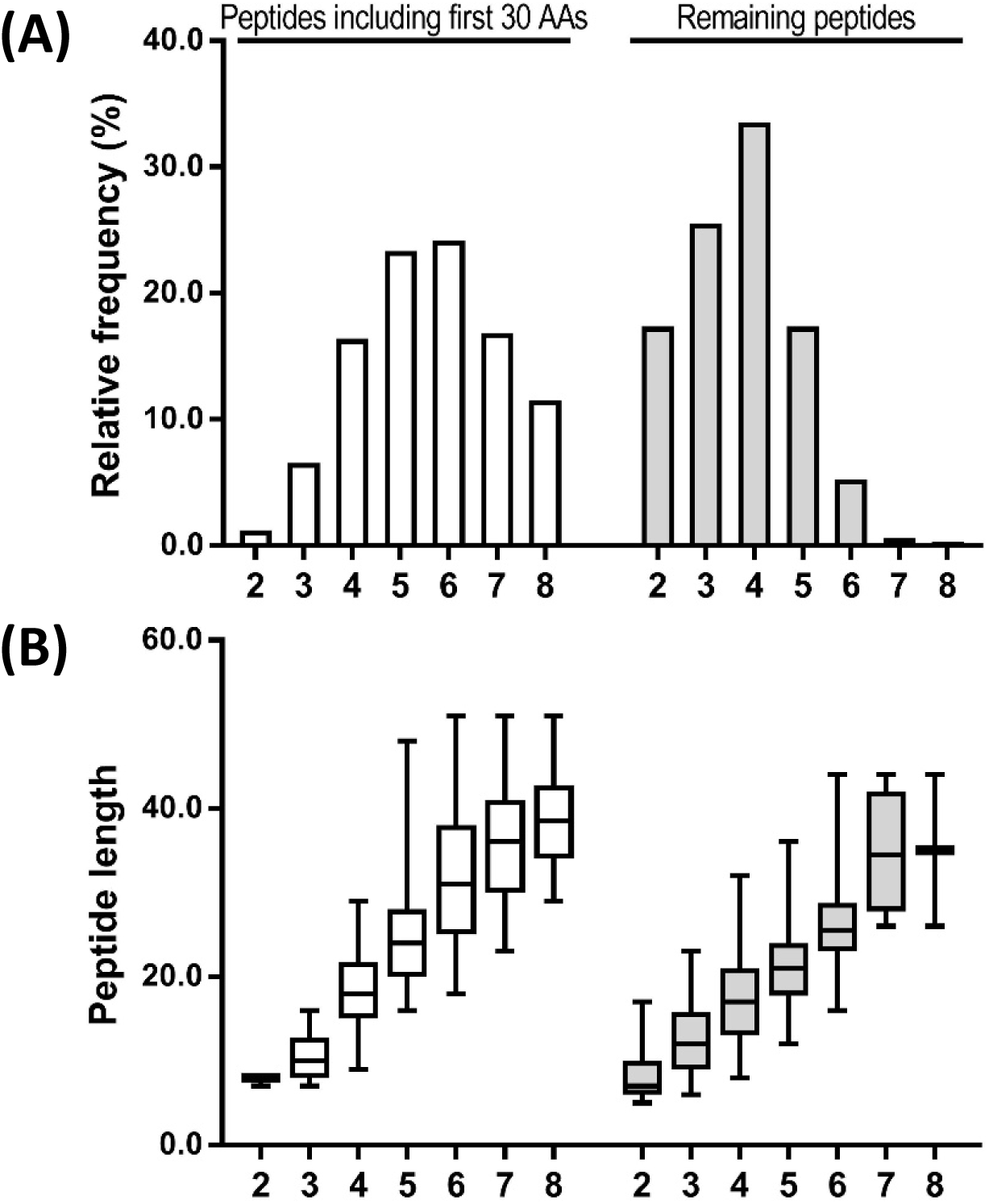
Properties of peptides identified within the tails (containing first 30 AAs) and within the remaining histone sequences by ETD. These include the charge (A) and length (B) Peptide length is shown in the form of a boxplot. Peptides identified in at least two replicates from each histone have been used in this analysis.

## References

1. Luger, K., Mäder, A.W., Richmond, R.K., Sargent, D.F., Richmond, T.J.: Crystal structure of the nucleosome core particle at 2.8 A resolution. Nature. 389, 251–260 (1997)

2. Strahl, B.D., Allis, C.D.: The language of covalent histone modifications. Nature. 403, 41–45 (2000)

3. Luger, K., Richmond, T.J.: The histone tails of the nucleosome. Curr. Opin. Genet. Dev. 8, 140–146 (1998)

4. Arents, G., Burlingame, R.W., Wang, B.C., Love, W.E., Moudrianakis, E.N.: The nucleosomal core histone octamer at 3.1 A resolution: a tripartite protein assembly and a left-handed superhelix. Proc. Natl. Acad. Sci. U. S. A. 88, 10148–10152 (1991)

5. Berman, H.M., Westbrook, J., Feng, Z., Gilliland, G., Bhat, T.N., Weissig, H., Shindyalov, I.N., Bourne, P.E.: The Protein Data Bank. Nucleic Acids Res. 28, 235–242 (2000)

6. Balasubramaniam, D., Komives, E.A.: Hydrogen-exchange mass spectrometry for the study of intrinsic disorder in proteins. Biochim. Biophys. Acta. 1834, 1202–1209 (2013)

7. Malvina Papanastasiou, James Mullahoo, Katherine C. DeRuff, Besnik Bajrami, Stephen E Johnston, Ryan Peckner, Samuel A Myers, Steven A Carr, Jacob D Jaffe: Chasing tails: Cathepsin-L improves structural analysis of histones by HX-MS. Molecular and Cellular Proteomics, in revision.

8. Schräder, C.U., Lee, L., Rey, M., Sarpe, V., Man, P., Sharma, S., Zabrouskov, V., Larsen, B., Schriemer, D.C.: Neprosin, a Selective Prolyl Endoprotease for Bottom-up Proteomics and Histone Mapping. Mol. Cell. Proteomics. 16, 1162–1171 (2017)

9. Hamuro, Y., Zhang, T.: High-Resolution HDX-MS of Cytochrome c Using Pepsin/Fungal Protease Type XIII Mixed Bed Column. J. Am. Soc. Mass Spectrom. 30, 227–234 (2019)

10. Nirudodhi, S.N., Sperry, J.B., Rouse, J.C., Carroll, J.A.: Application of Dual Protease Column for HDX-MS Analysis of Monoclonal Antibodies. J. Pharm. Sci. 106, 530–536 (2017)

11. Cravello, L., Lascoux, D., Forest, E.: Use of different proteases working in acidic conditions to improve sequence coverage and resolution in hydrogen/deuterium exchange of large proteins. Rapid Commun. Mass Spectrom. 17, 2387–2393 (2003)

12. Mazon, H., Marcillat, O., Forest, E., Vial, C.: Local dynamics measured by hydrogen/deuterium exchange and mass spectrometry of creatine kinase digested by two proteases. Biochimie. 87, 1101–1110 (2005)

13. Man, P., Montagner, C., Vernier, G., Dublet, B., Chenal, A., Forest, E., Forge, V.: Defining the interacting regions between apomyoglobin and lipid membrane by hydrogen/deuterium exchange coupled to mass spectrometry. J. Mol. Biol. 368, 464–472 (2007)

14. Zhang, H.-M., Kazazic, S., Schaub, T.M., Tipton, J.D., Emmett, M.R., Marshall, A.G.: Enhanced digestion efficiency, peptide ionization efficiency, and sequence resolution for protein hydrogen/deuterium exchange monitored by Fourier transform ion cyclotron resonance mass spectrometry. Anal. Chem. 80, 9034–9041 (2008)

15. Zhang, H.-M., McLoughlin, S.M., Frausto, S.D., Tang, H., Emmett, M.R., Marshall, A.G.: Simultaneous reduction and digestion of proteins with disulfide bonds for hydrogen/deuterium exchange monitored by mass spectrometry. Anal. Chem. 82, 1450–1454 (2010)

16. Englander, J.J., Del Mar, C., Li, W., Englander, S.W., Kim, J.S., Stranz, D.D., Hamuro, Y., Woods, V.L., Jr: Protein structure change studied by hydrogen-deuterium exchange, functional labeling, and mass spectrometry. Proc. Natl. Acad. Sci. U. S. A. 100, 7057–7062 (2003)

17. Mayne, L., Kan, Z.-Y., Chetty, P.S., Ricciuti, A., Walters, B.T., Englander, S.W.: Many overlapping peptides for protein hydrogen exchange experiments by the fragment separation-mass spectrometry method. J. Am. Soc. Mass Spectrom. 22, 1898–1905 (2011)

18. Rand, K.D., Adams, C.M., Zubarev, R.A., Jørgensen, T.J.D.: Electron capture dissociation proceeds with a low degree of intramolecular migration of peptide amide hydrogens. J. Am. Chem. Soc. 130, 1341–1349 (2008)

19. Syka, J.E.P., Coon, J.J., Schroeder, M.J., Shabanowitz, J., Hunt, D.F.: Peptide and protein sequence analysis by electron transfer dissociation mass spectrometry. Proc. Natl. Acad. Sci. U. S. A. 101, 9528–9533 (2004)

20. Guthals, A., Clauser, K.R., Frank, A.M., Bandeira, N.: Sequencing-grade de novo analysis of MS/MS triplets (CID/HCD/ETD) from overlapping peptides. J. Proteome Res. 12, 2846–2857 (2013)

21. Kan, Z.-Y., Walters, B.T., Mayne, L., Englander, S.W.: Protein hydrogen exchange at residue resolution by proteolytic fragmentation mass spectrometry analysis. Proc. Natl. Acad. Sci. U. S. A. 110, 16438–16443 (2013)

22. Karch, K.R., Coradin, M., Zandarashvili, L., Kan, Z.-Y., Gerace, M., Englander, S.W., Black, B.E., Garcia, B.A.: Hydrogen-Deuterium Exchange Coupled to Top- and Middle-Down Mass Spectrometry Reveals Histone Tail Dynamics before and after Nucleosome Assembly. Structure. (2018). doi: 10.1016/j.str.2018.08.006

